# DENND2B activates Rab35 at the intercellular bridge regulating cytokinetic abscission and tetraploidy

**DOI:** 10.1101/2023.01.12.523789

**Authors:** Rahul Kumar, Vincent Francis, Maria S. Ioannou, Adriana Aguila, Emily Banks, Gopinath Kulasekaran, Maleeha Khan, Peter S. McPherson

## Abstract

Cytokinesis is the final stage of cell division. Successful cytokinesis requires membrane trafficking pathways regulated by Rabs, molecular switches activated by guanine nucleotide exchange factors (GEFs). Late in cytokinesis, an intercellular cytokinetic bridge (ICB) connecting the two daughter cells undergoes abscission, which requires depolymerization of actin. Rab35 recruits MICAL1 to oxidate and depolymerize actin filaments. We report that DENND2B, a protein previously implicated in cancer, mental retardation and multiple congenital disorders functions as a GEF for Rab35 and recruits and activates the GTPase at the ICB. Unexpectedly, the N-terminal region of DENND2B interacts with an active mutant of Rab35, suggesting that DENND2B is both a Rab35 GEF and effector. Knockdown of DENND2B delays abscission resulting in increased multinucleated cells and overaccumulation of F-actin at the ICB. F-actin accumulation leads to formation of a chromatin bridge, a process known to activate the NoCut/abscission checkpoint, and DENND2B knockdown actives Aurora B kinase, a hallmark of checkpoint activation. This study identifies DENND2B as a crucial player in cytokinetic abscission and provides insight into the multisystem disorder associated with DENND2B mutation.

## Introduction

Membrane trafficking controls the localization and levels of proteins important for cell physiology and alterations in these pathways cause disease. Key to membrane trafficking pathways are small GTPases including Rabs [1], molecular switches that toggle between a guanosine diphosphate (GDP)–bound inactive state and a guanosine triphosphate (GTP)–bound active state [2]. Rabs are activated by guanine nucleotide exchange factors (GEFs) [3], the largest family being the DENN domain-bearing proteins that comprise at least 18 members, most of which are poorly characterized [4]. Upon activation, Rabs in their GTP-bound state bind to effectors, proteins that carry out downstream steps in membrane trafficking [5].

Cytokinesis, the final step of cell division requires delivery and retrieval of select proteins to the site of daughter cell separation [6][7]. Defects in cytokinesis are associated with cancers, Lowe syndrome and neurodevelopmental disorders [8][9][10][11][12][13][14]. Cytokinesis involves the physical separation of two daughter cells at the end of mitosis/meiosis. Before abscission, the daughter cells remain attached through an intercellular bridge (ICB), which can persist for several hours [15]. The ICB contains an array of cytoskeletal components functioning in the ingression of the plasma membrane, which results in a cytokinetic furrow [6][16]. Successful abscission requires clearance of cytoskeletal elements such as microtubules and filamentous actin (F-actin) allowing for constriction of the plasma membrane by the ESCRT machinery [17][18][19]. The removal of microtubules from the ICB depends on ESCRT-mediated delivery of the microtubule-depolymerizing enzyme Spastin [20][21]. F-actin clearance involves the small GTPase Rab35. Rab35 prevents accumulation of F-actin through its effector Oculo-Cerebro-Renal syndrome of Lowe, an inositol (4,5)P2 5-phosphatase [12][22], and drives F-actin depolymerization through the effector MICAL1, which oxidates and depolymerizes F-actin [23][24]. Rab35/MICAL1-dependent actin depolymerization is also important for the recruitment of ESCRT [23][24]. A scaffold protein, Rab11FIP1, gets recruited to the ICB after recruitment of Rab35 and helps maintain Rab35 at this site [25]. The activation and recruitment of Rab35 at the ICB remains poorly understood.

There are pathophysiological conditions such as high membrane tension, defective nuclear pore complex integrity, and the presence of trapped chromatin in the cleavage plane (chromatin bridge) that trigger the abscission (NoCut) checkpoint machinery, which delays completion of abscission and leads to cytokinetic failure, tetraploidy and the development of cancer [26][27][28][29][30][31][31][32][33][34]. The activation of the checkpoint machinery is also observed with accumulation of F-actin at the ICB [31][35][36], although a recent report suggested F-actin accumulation may not coincide with checkpoint activation [37].

As Rabs become more commonly linked to disease [38][39], there is increased interest in identifying and characterizing their activators (GEFs). The largest family of GEFs contain an evolutionary conserved protein module, the DENN (differentially expressed in normal and neoplastic cells) domain [4]. There are minimally 18 DENN-domain bearing proteins and most are poorly characterized [4]. An *in vitro* screen based on purified Rabs and 16 DENN domain proteins led to the assignment of a single, unique Rab to each DENN domain subfamily [40]. Subsequent cell biological studies focusing on individual DENN domain proteins such as DENND1C [41] and DENND2B [42] revealed different Rab substrates than those identified in the *in vitro* screen. This disparity could stem from the fact that *in vitro* GEF assays are challenging as purification of recombinant Rabs can lead to their inactivation and altered nucleotide loading [43], and inactivation of purified DENN domains may also occur due to misfolding.

We developed a cell-based assay to identify Rab substrates for DENN domain proteins [44]. The basis of this assay is the finding that GEFs are primarily responsible for driving the spatial and temporal localization of Rab GTPases [44]. For example, when GEFs are artificially targeted to organelles such as mitochondria, Rab substrates relocalize to these new sites [44]. Utilizing this approach, we discovered that DENN domain proteins activate a larger array of Rab substrates than previously thought [45]. Through investigation of newly identified DENN/Rab GEF/substrate pairs we discovered that DENND2B activates Rab10 to regulate primary cilia formation [45]. DENND2B is multifunctional in that it also acts as a GEF for Rab13, promoting the development of epithelial cancer [42]. Additionally, DENND2B functions in nervous system development and a loss of function mutation in DENND2B leads to severe mental retardation, seizures, bilateral sensorineural hearing loss, unilateral cystic kidney dysplasia, frequent infections and other congenital anomalies [46].

Here, we demonstrate that DENND2B interacts with Rab35 as both an effector and a GEF, and in so doing leads to enrichment of activated Rab35 at the ICB, allowing for actin depolymerization and cytokinesis. Disruption of DENND2B leads to increased formation of a chromatin bridge at the ICB and activates the NoCut/abscission checkpoint, delaying abscission. We suggest that these processes contribute to the role of DENND2B in cancer and neurodevelopmental disorders.

## Results

### Loss of DENND2B delays cytokinetic abscission

DENND2B is implicated in cancer and neurodevelopmental, processes involving cytokinetic defects [42][46][8][9][10][12][13][14]. We observed that loss of DENND2B causes cells to grow slowly and thus examined if DENND2B functions in cytokinesis. We performed phase contrast time-lapse microscopy imaging of HeLa cells comparing control and DENND2B knockdown (KD) using previously reported shRNA sequences [42][47]) that lead to a ~75% reduction in DENND2B mRNA levels (Fig. 1A). These shRNAs reduce DENND2B protein levels in human cells [47]. Both control and KD cells form the ICB following furrow ingression (Fig. 1B) but control cells take on average 237 min to complete cytokinesis (as previously reported for HeLa cells [23]) whereas DENND2B KD cells require 472 min (Fig. 1B-D), similar to the delay observed for depletion of MICAL1 depletion [12][23]. DENND2B KD cells also have an increase in binucleated cells (Fig. S1A-B), a phenotype observed upon KD of Rab35 or Rab11FIP1 [25][48]. Thus, DENND2B is a positive regulator of cytokinetic abscission.

**Figure 1.**
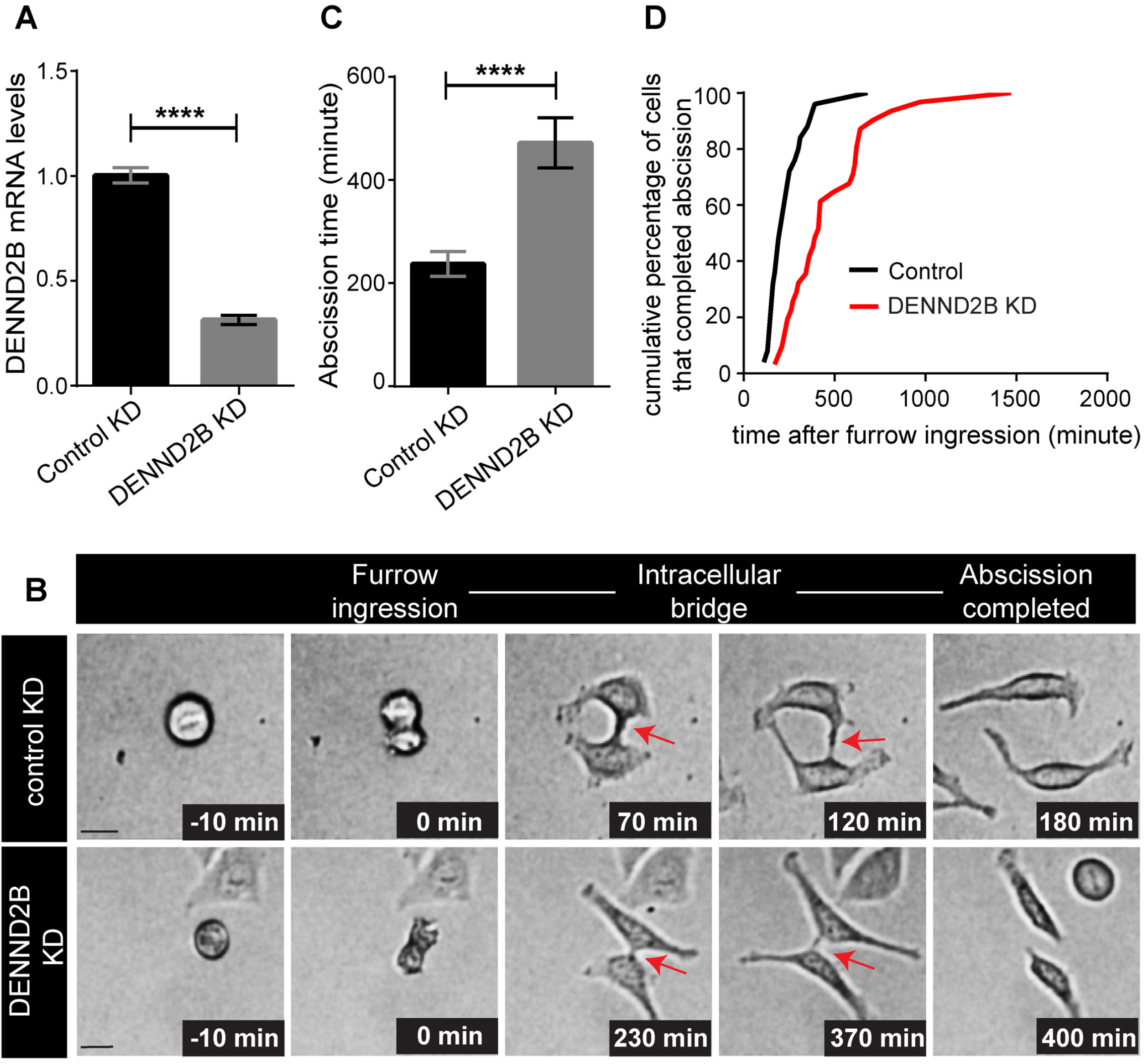
DENND2B is required for cytokinetic abscission. **(A)** HeLa cells were transduced with control or DENND2B shRNA lentivirus and DENND2B mRNA levels were quantified by real-time PCR; mean ± SEM; unpaired t-test (****, P ≤ 0.0001). **(B)** Representative images of time-lapse phase-contrast microscopy of control or DENND2B KD HeLa cells. Time zero is set as the frame of furrow ingression. Scale bars =20 μm. The red arrow represents the cytokinetic bridge. **(C)** Quantification of mean abscission timing of cytokinesis in control or DENDN2B KD cells; mean ± SEM; Mann-Whitney U test (****, P < 0.0001; >30 cells per condition). **(D)** Control or DENND2B KD HeLa cells were imaged by time-lapse microscopy. The plot represents the cumulative percentage of cells that completed abscission (abscission time).

### DENND2B regulates cytokinetic abscission via Rab35

A screen of the DENN domain of DENND2B against all 60 Rabs using a cell-based mitochondrial recruitment assay [45] revealed Rab8A, Rab8B, Rab10, Rab13, Rab15, Rab27A, Rab27B, and Rab35 as potential DENND2B GEF substrates [45]. Of these, Rab35 and Rab8 were identified in an unbiased proteomic analysis of the midbody, an organelle assembled at the center of the ICB [49][50], pointing our attention to these two Rab substrates. Expression of active mutants of Rab35 (Rab35 Q67L) but not Rab8 (Rab8 Q67L) in DENND2B KD cells rescued the cytokinetic abscission defect (Fig. 2A-B). Expressing a DENND2B construct resistant to DENND2B shRNA also rescues the cytokinetic abscission timing, supporting the specificity of DENND2B function in cytokinesis (Fig. 2A-B). Collectively, these data indicate that DENND2B controls cytokinetic abscission, likely through Rab35.

**Figure 2.**
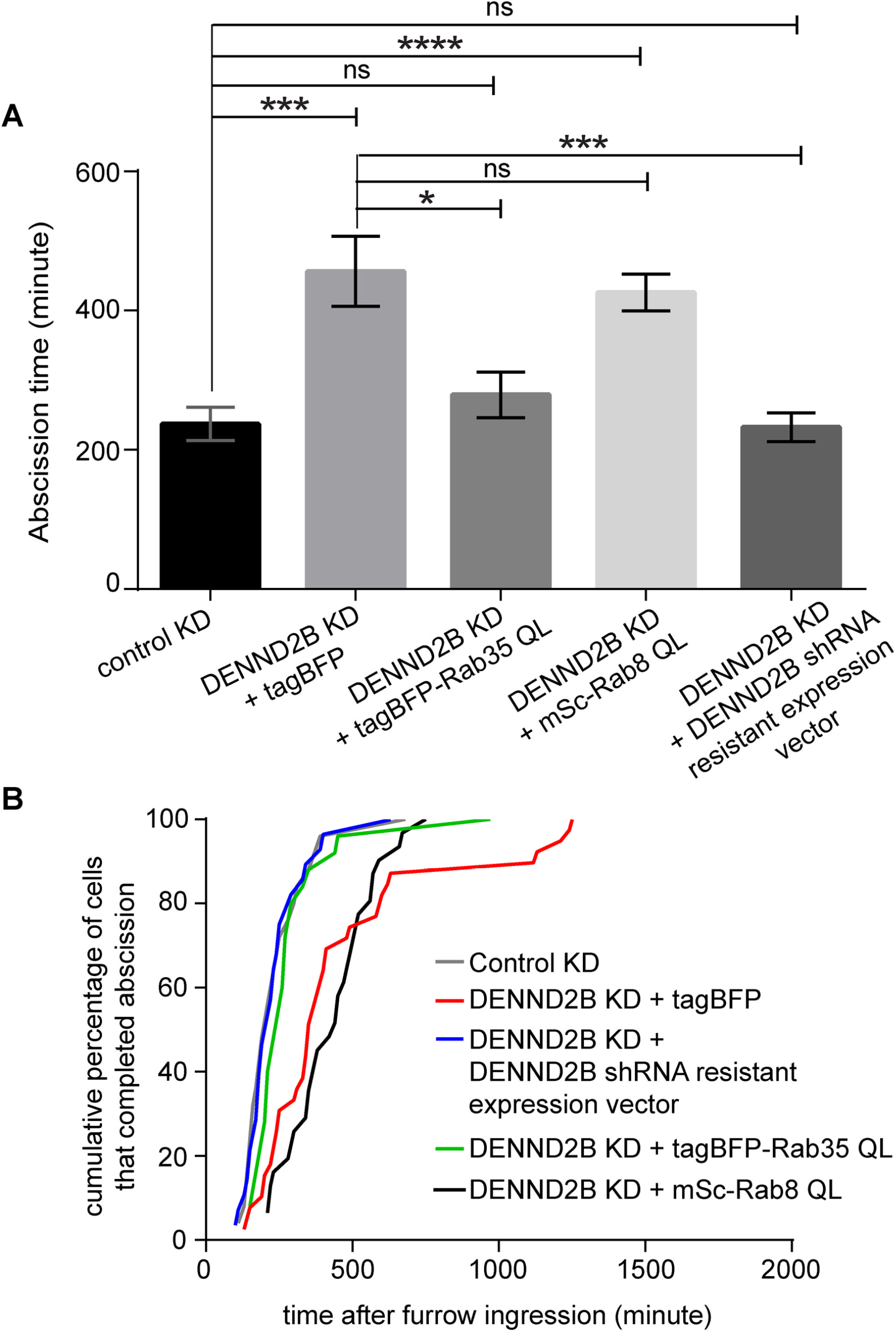
Expression of Rab35 active mutant in DENND2B KD cells rescues cytokinesis. **(A)** DENND2B KD HeLa cells were transduced with lentivirus mediating expression of tagBFP or tagBFP-Rab35 QL or mSc-Rab8 QL or mSc-DENND2B (resistant to DENND2B KD shRNA) and cytokinetic abscission timing was calculated using time-lapse phase contrast microscopy. The graph represents the quantification of mean abscission timing of cytokinesis; mean ± SEM; Kruskal-Wallis test, with pairwise multiple comparison test (****, P < 0.0001; ***, P < 0.001; **, P < 0.01; *, P < 0.05; >25 cells per condition) **(B)** HeLa cells in **A** were imaged by time-lapse microscopy. The data represents the cumulative percentage of cells that completed abscission (abscission time).

### DENND2B functions as a GEF for Rab35

Next, we sought to examine if DENND2B interacts with and/or serves as a GEF for Rab35. We next examined the potential GEF activity of DENND2B towards Rab35 using an assay in which GEFs can target substrates to the surface of mitochondria [44]. The mitochondrial targeted DENN domain of DENND2B [DENN(2B)-mito] leads to a near-complete steady-state re-localization of co-transfected GFP-Rab35 to mitochondria (Fig. 3A). While the mitochondrially targeted DENN domain of DENND2B induces a morphological change in the mitochondria, this is not relevant to the study as the assay aims simply to identify candidate GTPase relocalized to the mitochondria.

**Figure 3.**
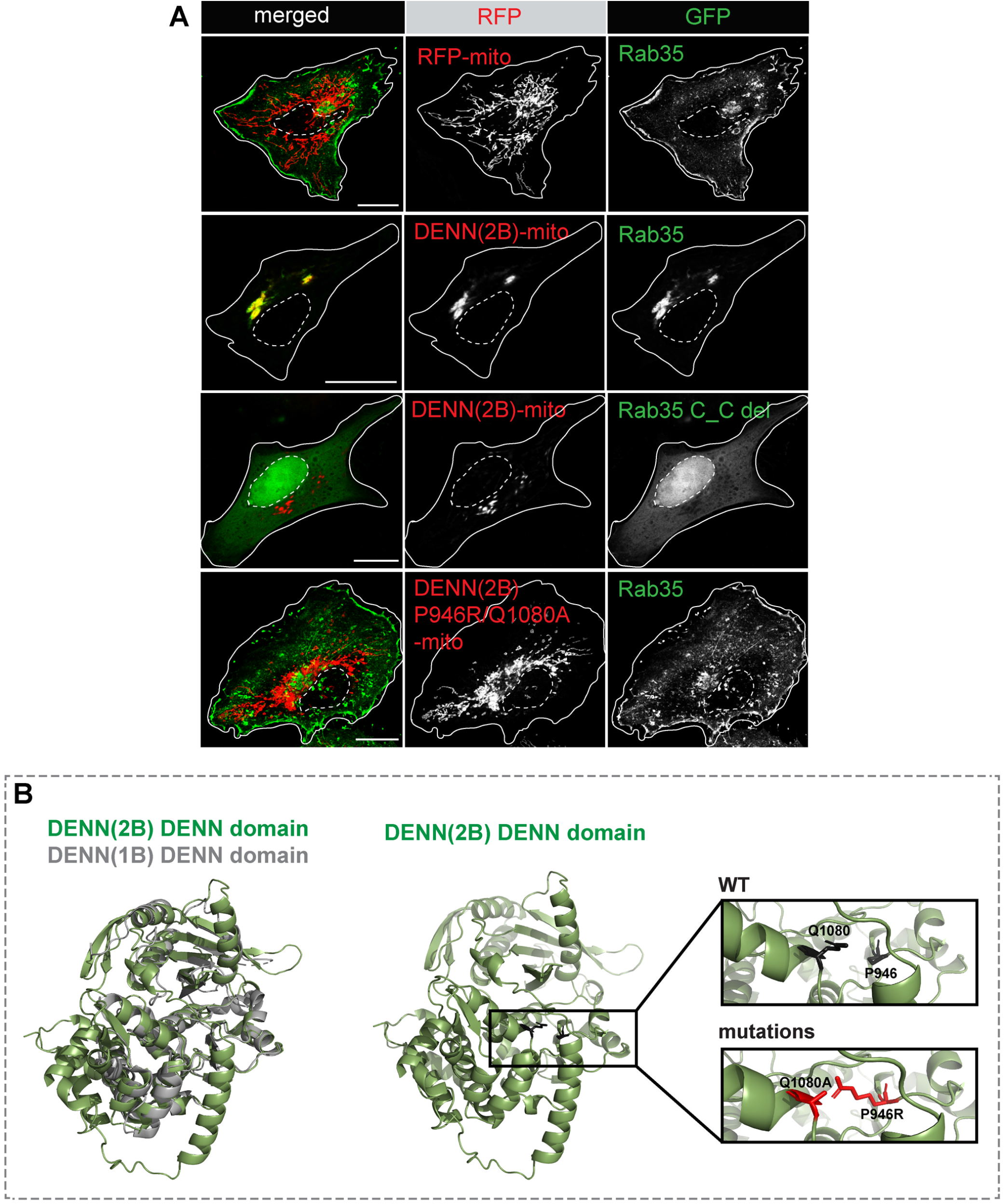
DENND2B functions as a GEF for Rab35. **(A)** HeLa cells co-transfected with GFP-Rab35 and RFP-mito or GFP-Rab35 and DENN(2B)-mito or DENN(2B)-mito and GFP-Rab35 C_C del or DENN(2B)-mito with double mutations P946R/Q1080A and GFP-Rab35. White solid line mark the cell periphery and dotted white line mark the nucleus. **(B)** Superposition of the crystal structure of the DENN domain of DENND1B [51] and an AlphaFold predicted structure of the DENN domain of DENND2B. Highlighted are two key residues involved in GEF activity and their mutation.

We next co-transfected HeLa cells with DENN(2B)-mito and a non-prenylatable form of Rab35 lacking the two C-terminal cysteines (GFP-Rab35 C_C del). GFP-Rab35 C_C del is not recruited to mitochondria (Fig. 3A), suggesting that Rab recruitment requires insertion of prenyl groups, a process mediated by GEF activity. Structural alignments of the DENN domain of DENND1B [51] with the DENN domain of DENND2B predicted by AlphaFold [52] (Fig. 3C) revealed that of the seven residues in the DENN domain of DENND1B important for interaction with and GEF activity towards Rab35 [51], all are conserved in DENND2B with three identical. Mutation of two identical residues (P946R and Q1080A) abolished the mitochondrial recruitment of Rab35 (Fig. 3B), reinforcing that the catalytic GEF activity of DENND2B is required for the activation of Rab35 and mitochondrial recruitment. Finally, we performed an effector binding assay using a glutathione S-transferase (GST) fusion with a C-terminal fragment of MICAL-L1 (molecule interacting with CasL-like 1), which selectively binds the active form of Rab35 [53]. The active levels of Rab35 were increased with DENND2B overexpression (Fig. 4A-B).

**Figure 4.**
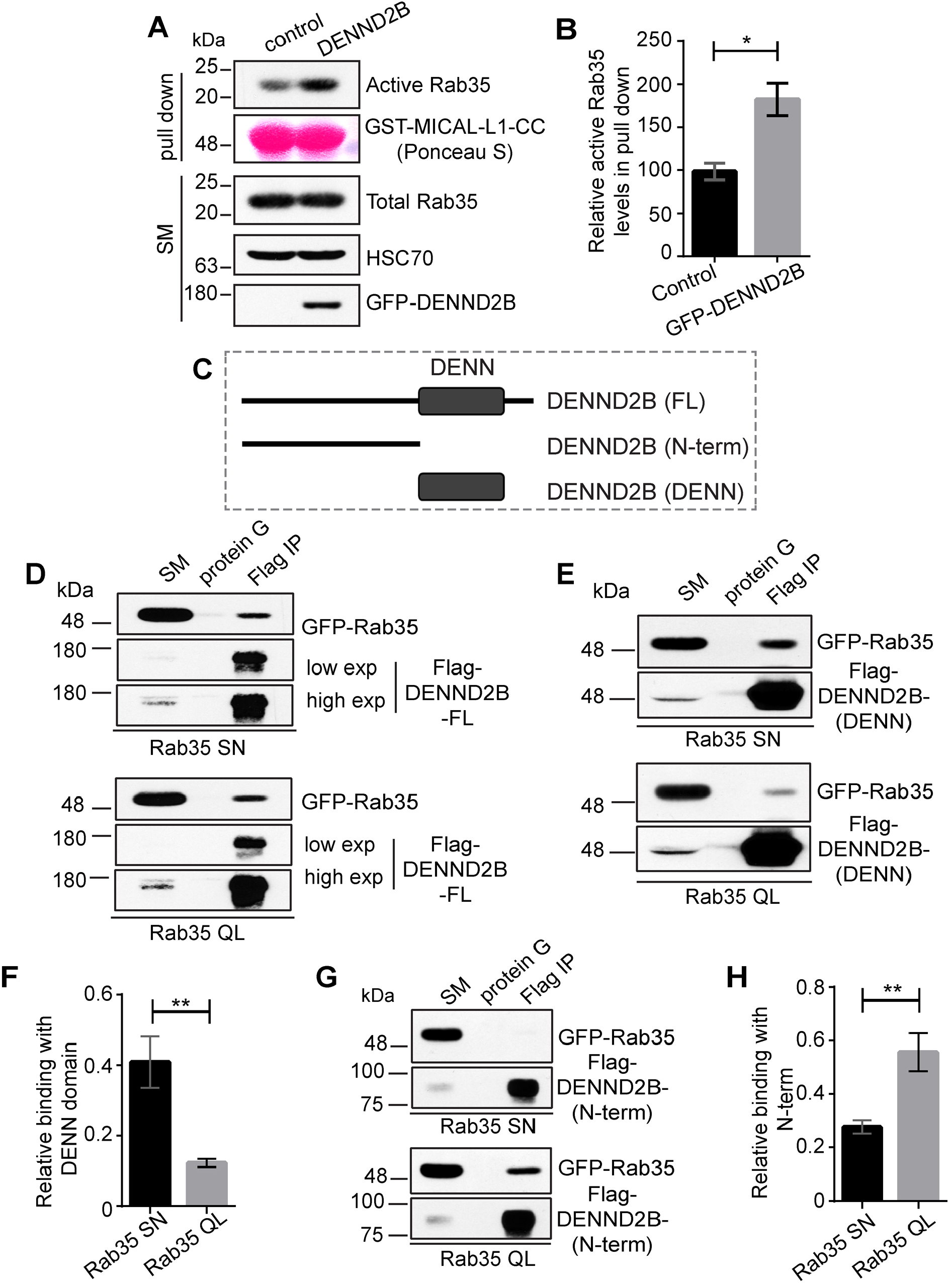
Nucleotide specificity of the DENN domain and the N-terminus of DENND2B interaction with Rab35. **(A)** HEK-293T cells were transfected with GFP-DENND2B. At 24 h post-transfection, transfected or untransfected (control) cell lysates were incubated with purified GST-MICAL-L1-CC. Specifically bound proteins were detected by immunoblot with anti-Rab35 antibody or anti-GFP antibody recognizing DENND2B or anti-HSC70 antibody for loading control. **(B)** Quantification of relative binding of active Rab35 with GST-MICAL-L1-CC from experiments as in **G**; mean ± SEM; unpaired t-test (*, P < 0.05; n = 3). **(C)** Schematic representation of various DENND2B constructs used in the biochemical experiments. **(D)** HEK-293T cells were co-transfected with Flag-DENND2B FL and GFP-Rab35 SN or GFP-Rab35 QL. At 24 h post-transfection, cells were lysed and incubated with protein G-agarose alone (mock) or protein G-agarose with anti-Flag antibody (Flag IP). Specifically bound proteins were detected by immunoblot with anti-GFP antibody to detect active/inactive Rab35 or anti-Flag antibody recognizing DENND2B. **(E)** HEK-293T cells were co-transfected with Flag-DENND2B (DENN) and GFP-Rab35 SN or GFP-Rab35 QL. At 24 h post-transfection, cells were lysed and incubated with protein G-agarose alone (mock) or protein G-agarose with anti-Flag antibody (Flag IP). Specifically bound proteins were detected by immunoblot with anti-GFP antibody to detect active/inactive Rab35 or anti-Flag antibody recognizing DENND2B (DENN). **(F)** Quantification of experiments as in **E**; mean ± SEM; unpaired t-test (**, P ≤ 0.01; n = 4). **(F)** HEK-293T cells were co-transfected with Flag-DENND2B (N-term) and GFP-Rab35 SN or GFP-Rab35 QL. At 24 h post-transfection, cells were lysed and incubated with protein G-agarose alone (mock) or protein G-agarose with anti-Flag antibody (Flag IP). Specifically bound proteins were detected by immunoblot with anti-GFP antibody to detect active/inactive Rab35 or anti-Flag antibody recognizing DENND2B (N-term). **(G)** Quantification of experiment as in **F**; mean ± SEM; unpaired t-test (**, P ≤ 0.01; n = 4).

A hallmark of GEF/Rab substrate relationships is that GEFs preferentially interact with the inactive, GDP-bound form of the Rab [51][54][55][56]. In coimmunoprecipitation experiments, full length (FL) Flag-DENND2B interacts with Rab35 but with no preference for active (QL) versus inactive (SN) mutants (Fig. 4C-D). In contrast, the DENN domain alone prefers inactive Rab35 whereas the N-terminal region (Fig. 4C) prefers the active form (QL) (Fig. 4E-H). Thus, through the N-terminal region, DENND2B appears to be an effector of Rab35. These data indicate that DENND2B is both a GEF for Rab35 and an effector of the same GTPase. We speculate that these dual interactions could serve a positive feedback role allowing DENND2B to rapidly, and strongly, activate Rab35 at specific cellular locations.

### DENND2B is localized in part to the midbody where it controls Rab35 recruitment and F-actin levels

We next investigated the localization of DENND2B in dividing cells. We expressed GFP-DENND2B in HeLa cells and stained with SiR-tubulin to visualize the progression of the ICB until complete scission [17][18]. Cleavage of the ICB is characterized by narrowing of either side of the midbody bulge (an organelle assembled at the center of the ICB) followed by scission, leaving behind a midbody remnant [17][18]. Upon furrow ingression, DENND2B accumulates at the cell-cell interface or the interface of the daughter cell and the ICB on either side (Fig. 5A). With the progression of the ICB, DENND2B accumulates at the cell-cell interface as revealed by the quantification of DENND2B fluorescence intensity from the marked representative regions (1 & 2) (Fig. 5A-B). Additionally, we detected a small fraction of DENNDB at the midbody as represented by the overlap of DENND2B and SiR-tubulin (Fig. 5A, C). The DENND2B intensity decreases at the cell-cell interface just before abscission (Fig. 5A-B). The midbody is an extremely dynamic structure. Therefore, we performed live-cell imaging on cells that were already dividing, and we were able to capture the midbody through multiple planes (Fig. 5D). DENND2B is present at the midbody throughout cytokinesis, even post cytokinetic abscission when the midbody has become a midbody remnant (Fig. 5D).

**Figure 5.**
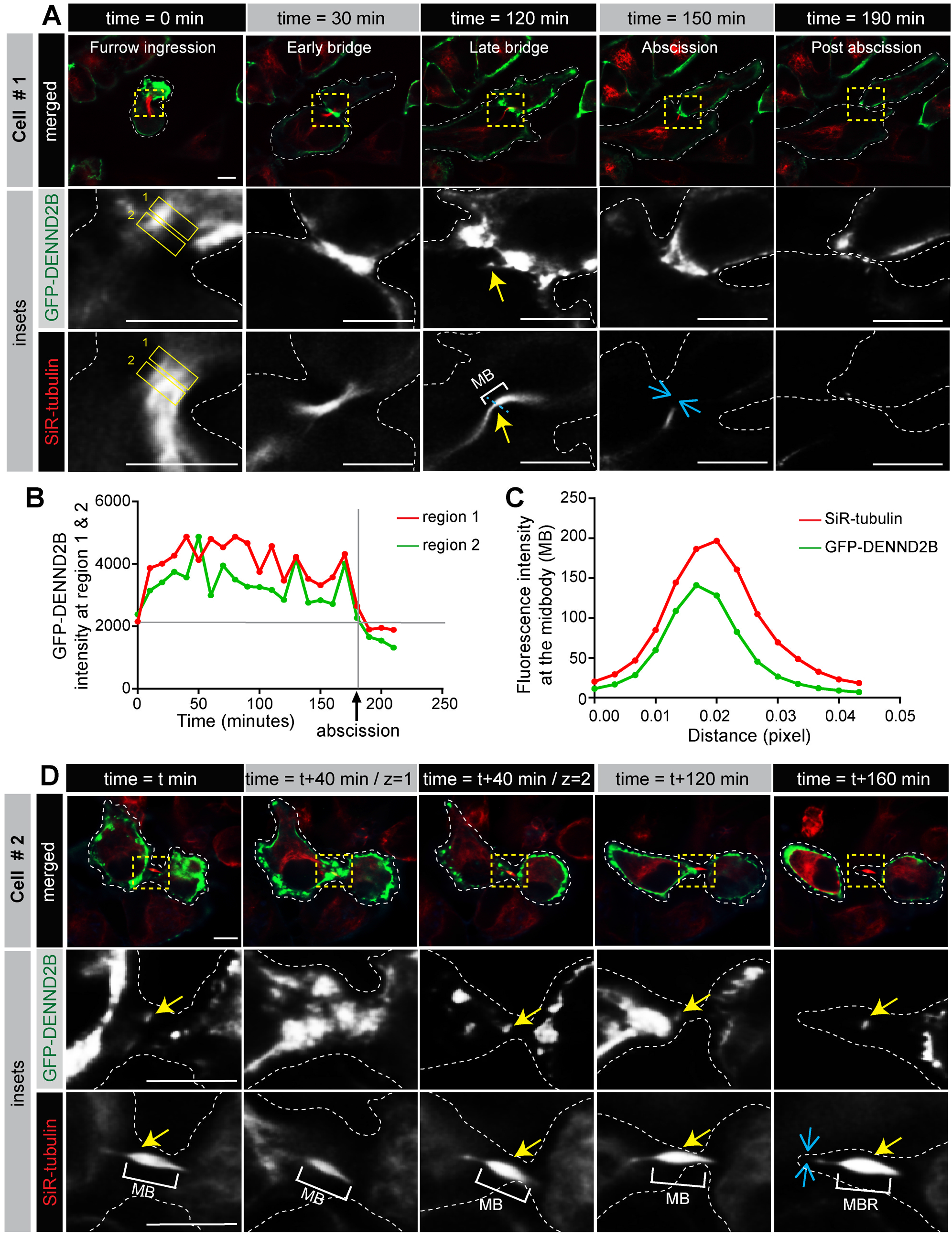
Time course of DENND2B recruitment at the cell-cell interface and midbody. **(A)** HeLa cells were transfected with GFP-DENND2B. At 24 h post transfection, cells were stained with SiR-tubulin and time-lapse imaging was performed using widefield fluorescence microscopy followed by deconvolution. Images represent frames of progressing cytokinesis over time, showing localization of DENND2B at the cell-cell interface and midbody. The yellow arrow represents localization of DENND2B at the midbody. The blue arrow represents completed abscission. Scale bars = 10 μm for the low-magnification images and 10 μm for the higher magnification insets (represented with yellow dotted box). The dotted white line marks the periphery of the dividing cell. **(B)** Plot of the fluorescence intensity of DENDN2B from the corresponding regions 1 and 2 drawn in **A** at the cell-cell interfaces throughout multiple frames over time. **(C)** Fluorescence intensity profiles along the dotted blue line from the inset image in **A** across the midbody (MB). **(D)** HeLa cells were processed as in **A** and already dividing cell was imaged over time and across multiple z-planes using widefield fluorescence microscopy followed by deconvolution. Images represent frames of progressing cytokinesis over time, showing localization of DENND2B at the cell-cell interface and midbody. The yellow arrow represents the localization of DENND2B at the midbody. The blue arrows represent completed abscission. Scale bars = 10 μm for the low-magnification images and 10 μm for the higher magnification insets (represented with yellow dotted box). Dotted white line marks the periphery of dividing cell.

GFP-DENND2B shows a similar distribution as that of tagBFP-Rab35 at the cell-cell interface at the late ICB (Fig. S2A-B). The enrichment pattern of Rab35 at the diving cell interface was the same as that previously reported [25]. Co-localization at the bridge supports that DENND2B and Rab35 function together in cytokinesis. Finally, it is known that GEFs control the localization of their Rab substrates [57]. To analyze the localization of Rab35 in the absence of DENND2B, we performed DENND2B KD in HeLa cells stably expressing tagBFP-Rab35 and stained with β-tubulin to identify dividing cells with a late bridge [23]. Cells showing Rab35 enrichment at cell/ICB interface of the late bridge decreased significantly in the DENND2B KD cells as compared to the control (Fig. 6A-B). Quantitative analysis of fluorescence intensity of Rab35 at the bridge revealed a ~2.5-fold decrease of Rab35 recruitment in the DENND2B KD cells (Fig. 6A, C).

**Figure 6.**
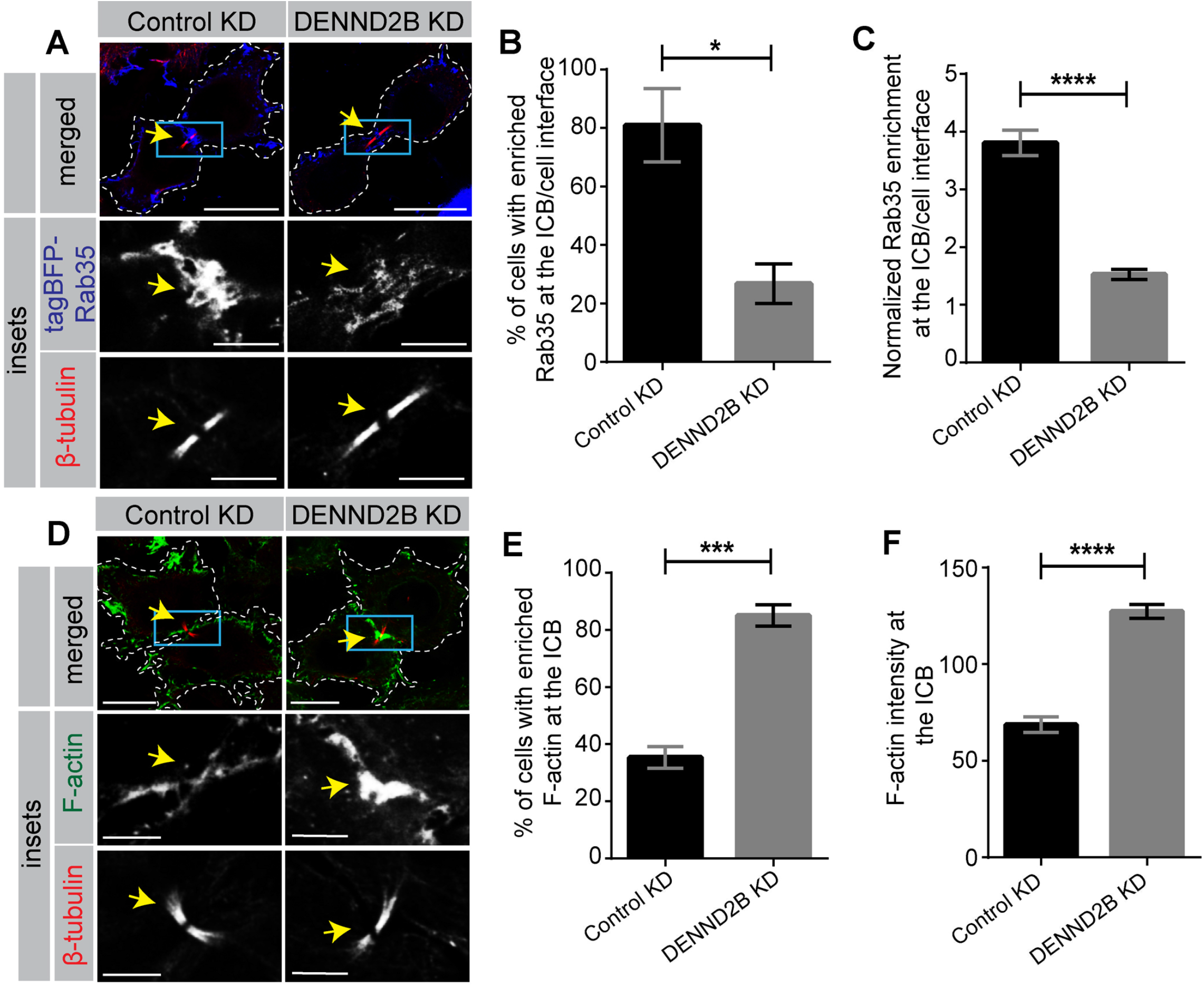
DENND2B KD causes loss of Rab35 and increased accumulation of F-actin at the cytokinetic bridge. **(A)** HeLa cells stably expressing tagBFP-Rab35 were transduced with control or DENND2B shRNA lentivirus. Cells were fixed and stained with β-tubulin to identify cytokinetic bridge. Scale bars = 20 μm for the low-magnification images and, 5.93 μm and 5.78 μm for the higher magnification insets corresponding to control or DENND2B KD cells (represented with blue box). The dotted white line marks the periphery of dividing cells. The yellow arrows represent cytokinetic bridges. **(B)** Quantification of experiment as in **A**; mean ± SEM; unpaired t-test (*, P ≤ 0.05; >25 cells per condition) **(C)** Quantification of experiments as in **A**; mean ± SEM; Mann-Whitney U test (****, P ≤ 0.0001; >25 cells per condition) **(D)** HeLa cells were transduced with control or DENND2B shRNA lentivirus. Cells were fixed and stained with Phalloidin for F-actin and β-tubulin. Scale bars = 20 μm for the low-magnification images and 7 μm for the higher magnification insets (represented with blue box). Dotted white line marks the periphery of dividing cell. The yellow arrows represent cytokinetic bridges. **(E)** Quantification of experiment as in **D**; mean ± SEM; unpaired t-test (***, P ≤ 0.001; >25 cells per condition) **(F)** Quantification of experiments as in **D**; mean ± SEM; unpaired t-test (****, P ≤ 0.0001; >25 cells per condition).

An important function of Rab35 at the ICB is to recruit MICAL1 to oxidate and depolymerize F-actin, facilitating cytokinetic abscission [23]. Since Rab35 recruitment at the late bridge is lost in the absence of DENND2B, we predicted that DENND2B depletion would alter F-actin levels at the bridge. There is an increase in the number of cells accumulating F-actin at the late bridge following DENND2B depletion (Fig. 6D-E) with a ~ 2-fold increase in F-actin levels within the bridge in DENDN2B KD cells (Fig. 6D, F). These data suggest that DENND2B activates Rab35 at the cytokinetic bridge with activated Rab35 driving downstream pathways causing F-actin depolymerization, facilitating cytokinetic abscission.

### DENND2B is recruited in the presence of lagging chromatin and regulates the abscission checkpoint machinery

Cytokinetic abscission is a tightly coordinated process, the timing of which is surveilled by a conserved complex of proteins known as the abscission checkpoint machinery [30]. The checkpoint machinery is activated when dividing cells are experiencing stress such as the presence of trapped chromatin (chromatin bridge) at the ICB, high membrane tension, or nuclear pore defects [30]. These stresses can alter cytokinesis and lead to tetraploidy [31].

Activation of the abscission checkpoint correlates with the accumulation of F-actin at the ICB [31][35][36][58] although accumulation of F-actin at the ICB is not always a consequence of this process [37]. Given that DENND2B KD leads to F-actin accumulation at the ICB, we sought to test if DENND2B functions in abscission checkpoint. In HeLa cells expressing GFP-DENND2B and stained with LAP2-β (a nuclear envelope protein) to observe chromatin bridges [31][35], we found an enriched pool of DENND2B at the midbody region with the LAP2-β positive chromatin bridge (Fig. 7A) and that an increased percentage of DENND2B depleted cells had accumulation of F-actin at the chromatin bridge as compared to other cellular locations (Fig. 7B-C). Fluorescence intensity quantification revealed a ~2-fold increase of Factin levels within the chromatin bridge in DENDN2B KD cells (Fig. 7B, D).

**Figure 7.**
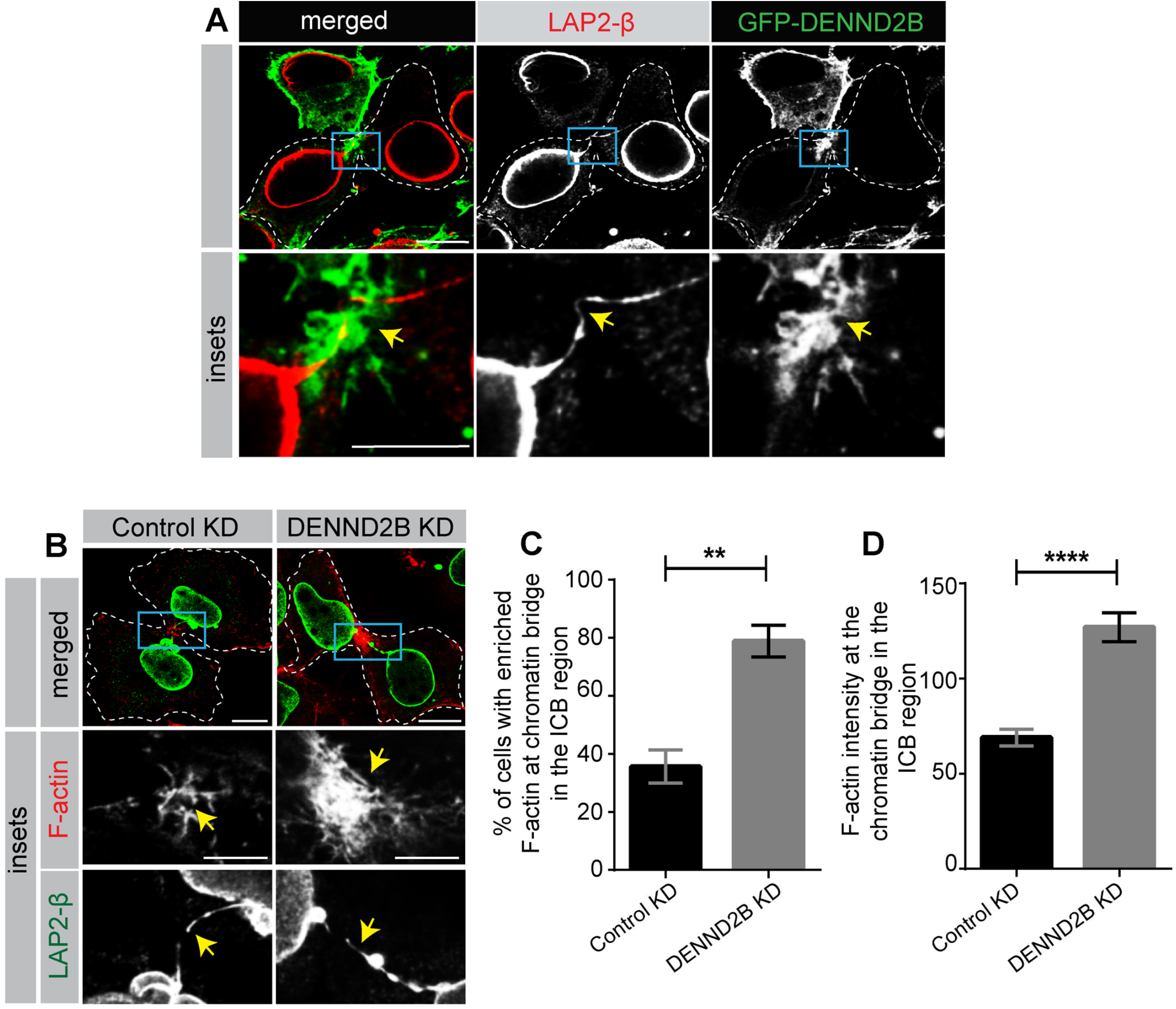
DENND2B is recruited to the midbody in the presence of chromatin bridges and controls F-actin levels. **(A)** HeLa cells were transduced with lentivirus mediating expression of GFP-DENDN2B. Cells were fixed and stained with LAP2-β. Scale bars = 12.7 μm for the low-magnification images and 6.35 μm for the higher magnification insets (represented with blue box). Dotted white line marks the periphery of dividing cell. **(B)** HeLa cells were transduced with control or DENND2B shRNA lentivirus. Cells were fixed and stained with Phalloidin for F-actin and LAP2-β for chromatin bridge. Scale bars = 14.3 μm and 14.8 μm for the low-magnification images corresponding to control and DENND2B KD cells and 7.15 μm and 7.4 μm for the higher magnification insets corresponding to control and DENND2B KD cells (represented with blue box). Dotted white lines mark the periphery of dividing cell. **(C)** Quantification of experiment in (B); mean ± SEM; unpaired t-test (**, P ≤ 0.01; >25 cells per condition) **(D)** Quantification of experiment in (B); mean ± SEM; unpaired t-test (****, *P* ≤ 0.01; >25 cells per condition).

To further demonstrate that accumulation of F-actin at the bridge is due to a lack of Rab35 recruitment, we expressed tagBFP-Rab35 QL (active mutant) or tagBFP alone in DENND2B KD cells and analyzed dividing cells with or without chromatin bridges. We found that the Rab35 active mutant localized at cytokinetic bridges (Fig. S3A, D). The percentage of DENND2B KD cells with accumulation of high F-actin at the bridge significantly decreased with the expression of Rab35 active mutant (Fig. S3B, E), and the F-actin intensities at the cytokinetic bridges were decreased by more than 2-fold in the presence of Rab35 active mutant, again suggesting that the activation and recruitment of Rab35 by DENND2B at the bridge with or without the chromatin is crucial for the maintenance of F-actin levels.

Lastly, we examined if DENND2B at the chromatin bridge regulates the abscission checkpoint. Aurora B kinase is a key regulator of abscission checkpoint that senses lagging chromatin and delays abscission [30][31][33], which is dependent on the activating phosphorylation of Aurora B (pT232 Aurora B) [30][59]. We observed that a pool of DENND2B colocalizes with active, phosphorylated Aurora B at the midbody upon activation of checkpoint by the lagging chromatin (Fig. 8A). A hallmark of the activation of the abscission checkpoint is the concentration of phospho-Aurora B at the midbody in the presence of a chromatin bridge [31]. Indeed, we found that the percentage of cells with phospho-Aurora B at the midbody of the chromatin bridge increased significantly (~2-fold) in the DENND2B KD cells as compared to the control (Fig. 8B-C), suggesting DENDN2B and Aurora B are functionally related. Finally, we also found that the percentage of cytokinetic bridges with the chromatin bridges also increased in the DENND2B KD cells as compared to the control (Fig. 8D-E), another hallmark of activation of checkpoint [37]. In summary, these data reveal that DENND2B also regulates abscission checkpoint and may function in the same pathway as Aurora-B.

**Figure 8.**
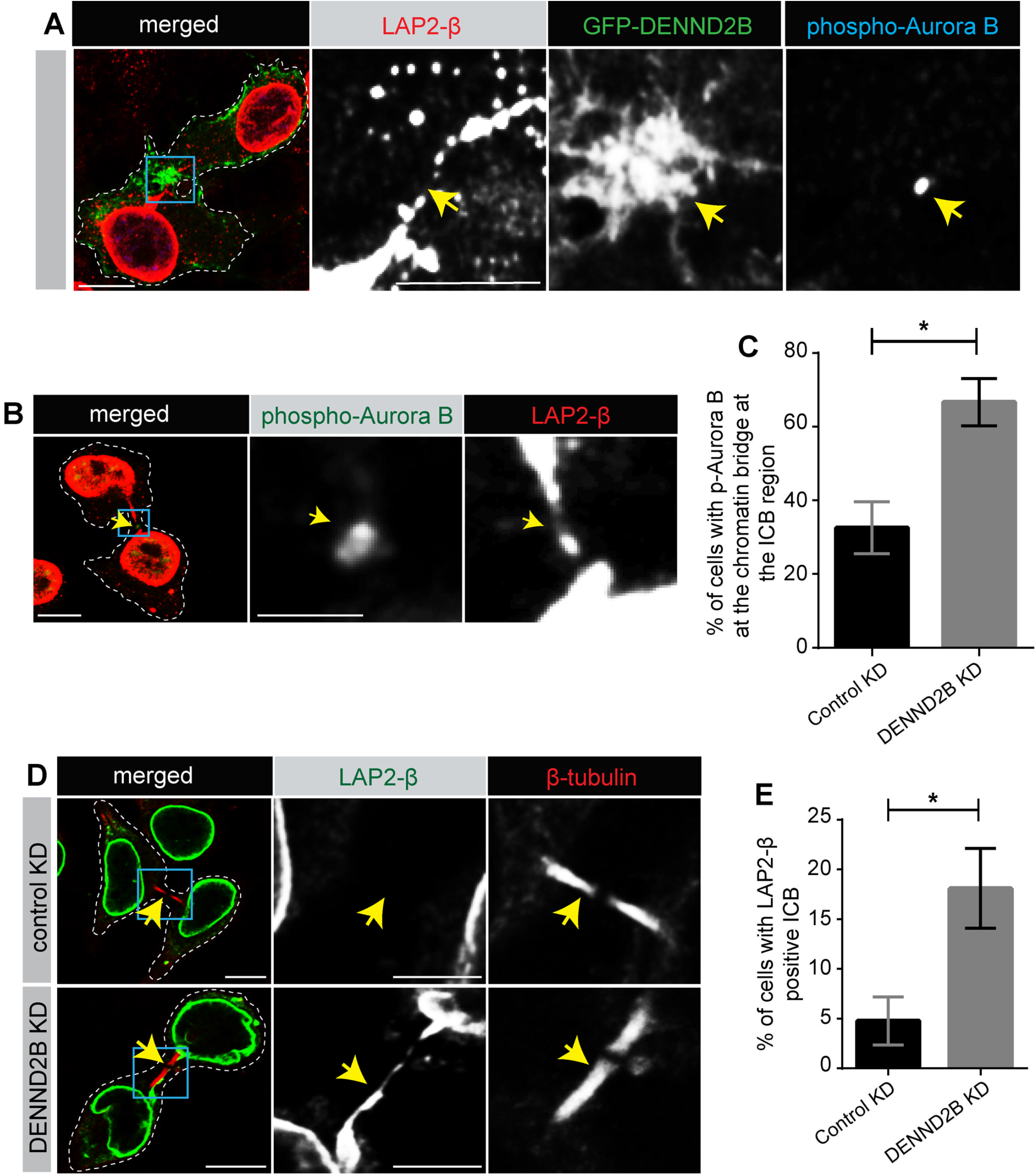
DENND2B colocalizes with Aurora B and functionally regulate abscission checkpoint. **(A)** HeLa cells were transduced with lentivirus mediating expression of GFP-DENDN2B, Cells were fixed and stained with LAP2-β and phospho-Aurora B. Scale bars = 10 μm for the low-magnification images and 5 μm for the higher magnification insets (represented with blue box). Dotted white line marks the periphery of dividing cell. Yellow arrow represents overlap of DENND2B, and phospho-Aurora B. **(B)** HeLa cells were fixed and stained with LAP2-β and phospho-Aurora B. Scale bars = 10 μm for the low-magnification images and 2.5 μm for the higher magnification insets (represented with blue box). Dotted white line marks the periphery of dividing cell. Yellow arrow represents presence of phospho-Aurora B at the lagging chromatin. **(C)** Quantification of experiment in (B); mean ± SEM; unpaired t-test (*, P ≤ 0.05; >25 cells per condition)**. (D)** HeLa cells were transduced with control or DENND2B shRNA lentivirus. Cells were fixed and stained with β-tubulin and LAP2-β. Scale bars = 10 μm for the low-magnification images and 5 μm for the higher magnification insets (represented with blue box). Dotted white line marks the periphery of dividing cell. Yellow arrow represents presence or absence of lagging chromatin at the bridge marked by β-tubulin. **(E)** Quantification of experiment in (D); mean ± SEM; unpaired t-test (*, P ≤ 0.05; >25 cells per condition).

## Discussion

Humans have ~60 Rabs and there is increasing interest to identify their activators given the growing association of altered membrane trafficking with human disease [60]. We identified multiple Rab GTPases as partners/substrates for DENN domain proteins [45]. We now report an important role of DENND2B in cytokinetic abscission.

Rab35 functions in various cellular contexts including cytokinesis [48][12][61], and yet, the mechanism by which Rab35 is activated and recruited at the cytokinetic bridge remains unknown. Here we demonstrate that DENND2B functions as a GEF for Rab35. Depletion of DENND2B leads to a delay in cytokinetic abscission and an increased number of binucleated cells [12][48]. F-actin clearance is required for the recruitment of ESCRT-III to allow timely abscission [23]. We observed accumulation of F-actin at the ICB upon loss of DENND2B. The idea that such phenotypes were caused by lost recruitment of Rab35 was validated by two findings. First, we observed a drastic reduction of cells with Rab35 at the cytokinetic bridge following DENND2B KD, and second, we could rescue the phenotype of F-actin accumulation at the bridge by expressing an active Rab35 mutant. Further, DENND2B and Rab35 colocalize at the cytokinetic bridge and show a similar enrichment pattern. Finally, we demonstrate that the DENN domain of DENND2B interacts with Rab35 in a nucleotide-dependent manner and that expression of DENND2B causes an increase in the levels of active Rab35 within the cell, further suggesting that DENND2B activates and recruits Rab35 at the cytokinetic bridge.

In addition to functioning as a GEF for Rab35, we find that DENND2B is a Rab35 effector. While effectors are recruited by Rab GTPases to help define the functional identity of the membrane [3], there are positive feedback loops caused by GEF–Rab-effector complexes [5]. These loops help maintain the membrane anchored GTPases and downstream signaling [5]. A well-known example of a GEF-Rab-effector complex is Rabex5/Rab5/Rabaptin5 [62]. Rabex5 has GEF activity for Rab5 upon initial recruitment of Rab5 to the membrane [62]. Once Rab5 is activated on the membrane, it interacts with its effector Rabaptin5 [63]. Subsequently, Rabaptin5 binds to Rabex5 and increases its GEF activity, thus ensuring sustained GTPase activation and effector function [64]. Another example of such a positive feedback mechanism involves polarized trafficking in yeast by the Sec2/Sec4/Sec15 complex. Activation of the Rab GTPase Sec4 is mediated by its exchange factor Sec2 and Sec2 binds to the effector of Sec4, Sec15 which could generate a positive feedback loop [3][65]. The fact that a portion of the same GEF (N-terminal fragment) is functioning as an effector as observed here for DENND2B raises an intriguing possibility that the N-terminal fragment could function to drive a positive feedback loop to help maintain Rab35 at the cytokinetic bridge, given that the bridge persists for up to several hours. Future experiments will seek to understand whether the N-terminus of DENND2B plays a role in a similar positive feedback mechanism to help sustain Rab35 remain anchored to the membrane and cause prolonged downstream effector function.

We also demonstrate DENND2B-mediated regulation of the abscission checkpoint. The absence of DENND2B increases the proportion of cells with cytokinetic bridges containing chromatin. This, together with the fact that DENND2B colocalizes with active phospho-Aurora B, a key component of the abscission checkpoint and that absence of DENND2B increases the number of cells with active phospho-Aurora B at the midbody of the bridge, indicates that DENND2B plays a role in regulating abscission checkpoint. Questions regarding the detailed mechanistic role of DENND2B in abscission checkpoint and potential DENND2B interaction with the checkpoint machineries remain unknown.

It is proposed that a balance of actin oxidation by MICAL1 and actin reduction by MsrB2 regulates F-actin levels at the cytokinetic bridge and controls abscission [35]. We propose that DENND2B activates Rab35 and recruits it to the cytokinetic bridge regardless of the presence of the chromatin bridge (Fig. 9). Recruited Rab35 contributes to F-actin oxidation mediated by MICAL1. In the absence of DENND2B, Rab35 recruitment is impaired resulting in elevated levels of F-actin at the bridge (Fig. 9). Accumulation of F-actin is not favorable for the recruitment of ESCRT-III, thus leading to delayed abscission [23]. The absence of DENND2B also activates abscission checkpoint in the presence of chromatin bridge (Fig. 9). A increased presence of activated checkpoint machinery (phospho-Aurora B) in the presence of chromatin at the bridge also contributes to the abscission delay [30] (Fig. 9).

**Figure 9.**
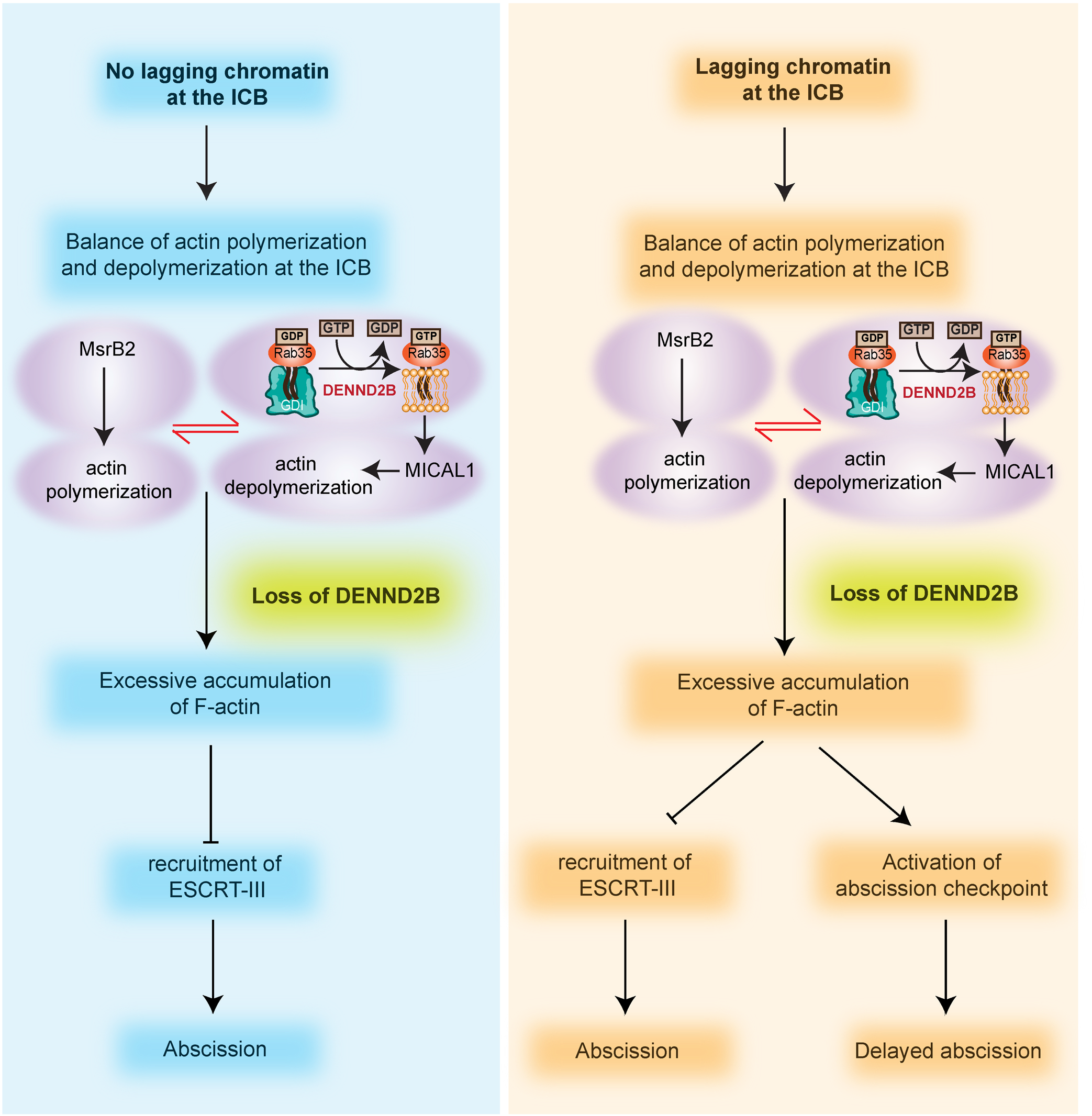
Proposed model of DENND2B regulation cytokinetic abscission. DENND2B activates and recruits Rab35 at the ICB with or without lagging chromatin. Active Rab35 recruits MICAL1 which mediates depolymerization of F-actin. In the absence of DENND2B, Factin accumulates at the bridge due to action of MsrB2 which polymerizes F-actin. F-actin accumulation inhibits ESCRT-III recruitment required for successful abscission. Whereas, in the presence of lagging chromatin, loss of DENND2B causes activation of abscission checkpoint which further contributes to delayed abscission.

With the identification of a new role of DENND2B/Rab35, we provide evidence supporting our previous findings that DENN domain proteins control a larger array of Rab GTPases in complex membrane trafficking pathways. Finally, this study leads to additional open questions. It appears that there are unknown contributing factors that define the context specific GEF activity of DENND2B for its activation at various cellular sites. While we have identified that DENND2B is crucial for the activation and recruitment of Rab35 at the bridge, the question regarding the upstream factors that cause the correct localization of DENND2B still needs to be determined.

In summary, we have uncovered that DENND2B functions as a GEF for Rab35 to control timely cytokinetic abscission and thereby prevents tetraploidy. We believe that the identification of this crucial pathway is a step forward in understanding the various congenital anomalies associated with a DENND2B loss-of-function patient mutation.

## Materials and Methods

### Cell lines

HeLa and HEK-293T cells were from ATCC (CCL-2 and CRL-1573)

### Cell culture

Cell lines were cultured in DMEM high-glucose (GE Healthcare cat# SH30081.01) containing 10% bovine calf serum (GE Healthcare cat# SH30072.03), 2 mM L-glutamate (Wisent cat# 609065), 100 IU penicillin and 100 μg/ml streptomycin (Wisent cat# 450201). Serum starvation media: DMEM high-glucose containing 2 mM L-glutamate, 100 IU penicillin and 100 μg/ml streptomycin. Cell lines were tested for mycoplasma contamination routinely using the mycoplasma detection kit (Lonza; cat# LT07-318).

### DNA constructs

RFP-mito, DENN(2B)-mito, DENN(2B) P946R/Q1080A-mito, GFP-Rab35, GFP-Rab35 C_C del, Flag-DENND2B (DENN) and Flag-DENND2B (N-term) were described previously [45][42]. The following constructs were generated by SynBio Technologies: mScarlet-DENND2B (in pmScarlet-i_C1), tagBFP-Rab35 (Human Rab35; in lentivirus vector pLVX-M-puro (Addgene 125839)), tagBFP (in pLVX-M-puro), mScarlet-Rab8 Q67L (Human Rab8a; in pLVX-M-puro) and mScarlet-DENND2B resistant to DENND2B shRNA (Xenopus DENND2B; in pLVX-M-puro). Lenti tagBFP-Rab35 Q67L construct was generated using the QuickChange lightning site-directed mutagenesis (SDM) kit (Agilent Technologies) on lentivirus vector pLVX-M-puro containing GFP-Rab35 using following primers: Fwd-5’ CACAGCGGGGCTGGAGCGCTTCC 3’ and Rev-5’ GGAAGCGCTCCAGCCCCGCTGTG 3’. Successful cloning of constructs was verified by sequencing.

### shRNA mediated knockdown of DENND2B

Production of control and DENND2B shRNA virus were described previously [42][47]. Briefly, the two shRNA sequences were used for control or DENND2B KD. The shRNA sequences were first cloned into pcDNA6.2/GW-emGFP–miR cassette and then the emGFP-miR cassette was PCR-amplified and subcloned into the pRRLsinPPT viral expression vector (Invitrogen). Control shRNA sequence: AATTCTCCGAACGTGTCACGT; DENND2B shRNA sequence:TGCTGCTTGGATGAAGCCAGCAAACAGTTTTGGCCACTGACTGACTGTTTGCTCTTCATCCAAG. The lentiviral particles were produced using shRNA containing pRRLsinPPT viral expression vector, pMD2.g, and pRSV-Rev as previously described [66].

### Real-time quantitative PCR

Total RNA was extracted from HeLa cells using RNeasy Mini kit (Qiagen) and 500 ng of RNA was used for the cDNA synthesis using iScript™ Reverse Transcription Supermix (Bio-Rad Laboratories). Real-time quantitative PCR was performed using the Bio-Rad CFX Connect Real-Time PCR Detection System with SsoFast™ EvaGreen Supermix (Bio-Rad Laboratories). The values were expressed as fold change in mRNA expression in cells relative to control WT cells (untreated) using TATA-box binding protein (TBP) and beta-2-microglobulin (B2M) as endogenous controls. The primer sequences used in this study were:

DENND2B-Fwd 5’ AGCAGAAAATCCTTTTGAGTTTG 3’,
DENND2B-Rev 5’ CTTTGGACAAGCTTGGGAATGC 3’.

### Protein expression using lentivirus

For each virus, HEK-293T cells (90% confluency) were transfected with 30 μg of lentivirus construct expressing protein of interest, 30 μg of psPAX2 and 15 μg of pMD2 VSV-G using linear polyethylenimine (PEI) [67], 25,000 Da (1mg/ml; Polysciences, Inc.). The transfection ratio was 1 μg plasmid:3 μL PEI. At 8 h post-transfection, culture media was replaced with collection media (15 ml per plate; regular medium supplemented with 1× nonessential amino acids and 1 mM sodium pyruvate). The media was collected at 24 and 36 h and replaced with fresh media (15 ml per plate) with each collection. The collected media at 24 h was stored at 4°C until the last collection. The collected culture media were then filtered through a 0.45 μm PES membrane and concentrated by centrifugation (16 hour at 6800 rpm), and the resulting pellets were resuspended in DMEM in 1/5,000 of the original volume. Concentrated viruses were aliquoted and stored at −80°C.

Following virus production, concentrated lentivirus was added to the cells with minimum culture media (for example, 1 ml media in a well of 6-well dish), and the media was replaced with fresh culture media the following day (16-20h). The expression of the target protein was verified by fluorescence under microscope.

### Antibodies and reagents

Mouse monoclonal Flag (M2) antibody was obtained from Sigma-Aldrich (F3165). Rabbit polyclonal GFP (A-6455) from Invitrogen, rat monoclonal HSC70 antibody (WB-1:10,000) is from Enzo (ADI-SPA-815-F). Phalloidin 647 (), Alexa Fluor 488 and 647-conjugated rabbit secondary antibodies are from Invitrogen. anti-Rab35 antibody (WB-1:1000) is from Abcam (ab152138), anti-β-tubulin antibody (IF-1:2000) is from Invitrogen (32-2600), anti-LAP2-β (rabbit; IF-1:300) antibody is from Proteintech (14651-1-AP), anti-LAP2-β (mouse, IF-1:500) antibody is from BD Biosciences (611000), phospho(T232)-Aurora B (IF-1:200) is from Rockland (600-401-677). SiR-Tubulin (Cytoskeleton, Inc., CY-SC002) was used at 100 nM.

### Imaging

#### Fixed cell imaging

HeLa cells following treatment as per the experimental condition were plated on poly-l-lysine coated coverslips. Cells were fixed with warm 4% paraformaldehyde for 10 min at 37°C, permeabilized for 5 min in 0.1% Triton X-100 and blocked for 1 h in 1% BSA in PBS (blocking buffer). Coverslips were incubated in blocking buffer containing diluted primary antibodies and incubated overnight at 4°C. Cells were washed 3 × 10 min with blocking buffer and incubated with corresponding Alexa Fluorophore-conjugated secondary antibodies diluted 1:1000 in blocking buffer for 1 h at room temperature. Cells were washed 3 × 10 min with blocking buffer and once with PBS. Coverslips were mounted on a microscopic slide using fluorescence mounting media (DAKO, Cat# S3023).

Imaging was performed using a Leica SP8 laser scanning confocal microscope. Image analysis was done using Image J. All the representative images in the figures were prepared for publication using Adobe Photoshop (adjusted contrast and applied 1-pixel Gaussian blur) and then assembled with Adobe Illustrator.

#### Phase contrast live cell microscopy

A Day before imaging, HeLa cells (treatment as per the experimental condition) were plated in a glass bottom 35mm MatTek dish (~35,000 cells per plate). 24-hour post cell seeding, MatTek dish containing cells were imaged using Zeiss live cell inverted microscope equilibrated in 5% CO2 and maintained at 37 °C. Timelapse phase contrast images were recorded every 10 min for ~30-hour using a 10X Air NA 0.45 objective (Zen software).

#### Widefield fluorescence Live cell imaging and deconvolution

A Day before imaging, HeLa cells (treatment as per the experimental condition) were plated in a glass bottom 35mm MatTek dish (~35,000 cells per plate) with media containing 100nM SiR-Tubulin. 24-hour post cell seeding, MatTek dish containing cells were imaged using Zeiss live cell inverted microscope equilibrated in 5% CO2 and maintained at 37 °C. Timelapse phase contrast images were recorded every 10 min for ~30-hour using a 63X Oil NA 1.4 objective (Zen software). The acquired live-cell images were processed with the fast iterative deconvolution module present in the Zen software to have increased resolution.

### Generation of stable cell line

HeLa cells were transduced with lentivirus mediating expression of tagBFP-Rab35. 48-hour post transduction, cells were treated with media containing 2 μg/ml puromycin. Transduced cells were maintained in puromycin containing media for 8 days, with changing media every alternate day. Following this, resistant cells were sorted using FACS to enrich heterogenous population of cells having low to medium expression of Rab35. Sorted cells expressing Rab35 were maintained in media containing 2 μg/ml puromycin for a week and subsequently, experiments were conducted using these cells in media containing 1 μg/ml puromycin throughout.

### Quantification of protein enrichment at the bridge

Each image frame was opened individually in ImageJ. Then a region of interest (ROI) was drawn using the custom region draw function either around the entire dividing cell using the outline of the cell periphery or the cell-cell interface. Subsequently, the average fluorescence signal was measured from both, and the fluorescence intensity of the entire dividing cell was used to normalize the intensity at the cell-cell interface.

### Protein purification

GST-MICAL-L1-CC protein was expressed in Escherichia coli BL21 (500 μM isopropyl β-d-1-thiogalactopyranoside; Wisent Bioproducts; at room temperature for 16 h) and purified using standard procedure in Tris buffer (20 mM Tris, pH 7.4, 100 mM NaCl, 10 mM MgCl2, and 1 mM dithiothreitol) supplemented with protease inhibitors (0.83 mM benzamidine, 0.20 mM phenylmethylsulfonyl fluoride, 0.5 mg/ml aprotinin and 0.5 mg/ml leupeptin).

### Transfection

HEK-293T cells were transfected using calcium phosphate. HeLa cells were transfected using jetPRIME Transfection Reagent (Polyplus) according to the manufacturer’s protocol.

### Biochemical assays

#### Co-immunoprecipitation

HEK-293T cells grown to 60% confluency in 15-cm dishes were transfected with Flag-tagged and/or GFP-tagged constructs. At 24 h post-transfection, upon confirming >90% transfected cells using fluorescence microscope, cells were gently washed with PBS, scraped into lysis buffer (20 mM HEPES, 100 mM NaCl, 0.5 mM dithiothreitol, 10 mM MgCl2, 1% Triton X-100 pH 7.4) supplemented with protease inhibitors (0.83 mM benzamidine, 0.20 mM phenylmethylsulfonyl fluoride, 0.5 mg/ml aprotinin and 0.5 mg/ml leupeptin), incubated for 20 min on a rocker at 4°C, and the lysates were centrifuged at 305,000 g for 15 min at 4°C. For Flag IP, supernatants were incubated with prewashed protein G beads–Sepharose beads (GE Healthcare) for 1.5 hour (preclearing step). Following preclearing, supernatants were incubated with protein G–Sepharose beads and the anti-Flag antibody for 2 h at 4°C. Beads coupled to the Flag antibody were washed three times with the same lysis buffer, eluted in SDS-PAGE sample buffer, resolved by SDS-PAGE, and processed for immunoblotting.

#### Effector pull-down assay

Cells were gently washed with PBS, lyzed in lysis buffer (20 mM HEPES, 100 mM NaCl, 15 mM MgCl2, 1% Triton X-100 pH 7.4) supplemented with protease inhibitors, incubated for 20 min on a rocker at 4°C, and the lysates were centrifuged at 305,000 g for 15 min at 4°C. For GST-pulldown experiments, supernatants were incubated with GST fusion proteins precoupled to glutathione–Sepharose beads for 1.5 h at 4°C. GST beads attached to the fusion proteins were washed three times with the same lysis buffer, eluted in SDS-PAGE sample buffer, resolved by SDS-PAGE, and processed for immunoblotting.

### Immunoblot

Cell lysates were run on large 10% polyacrylamide gels and transferred to nitrocellulose membranes. Proteins on the blots were visualized by Ponceau staining. Blots were then blocked with 5% milk in Tris-buffered saline with 0.1% Tween 20 (TBST) for 1 h followed by incubation with antibodies overnight at 4°C diluted in 5% milk in TBST. Next day, blots were washed with TBST three times, each 10 minutes. Then, the peroxidase-conjugated secondary antibody was incubated in a 1:5000 dilution in TBST with 5% milk for 1 h at room temperature followed by washes (3 times, 10 minutes each).

### Statistics

Graphs were prepared using GraphPad Prism software. All statistical tests were performed using SPSS. For all data, the normality test was performed before determining the appropriate statistical test. For normally distributed data, comparisons were made using either T-test or one-way ANOVA. For non-normally distributed data, comparisons were made using Mann-Whitney U test or Kruskal-Wallis test. All data are shown as the mean +/- SEM with P < 0.05 considered statistically significant.

## Acknowledgement

We thank Jacynthe Philie and Maryam Fotouhi for their excellent technical assistance. We acknowledge the Neuro Microscopy Imaging Centre and Advanced BioImaging Facility at the McGill University. This work was supported by a Canadian Institutes of Health Research Foundation Grant to PSM. RK is supported by a studentship from ALS Canada. VF was supported by a fellowship from the Fonds de recherche du Quebec – Sante (FRQS). GK was supported by FRQS and a Jeanne Timmins Costello Fellowship. PSM is a Distinguished James McGill Professor and a Fellow of the Royal Society of Canada.

## Figure legends

**Supplementary figure 1.**
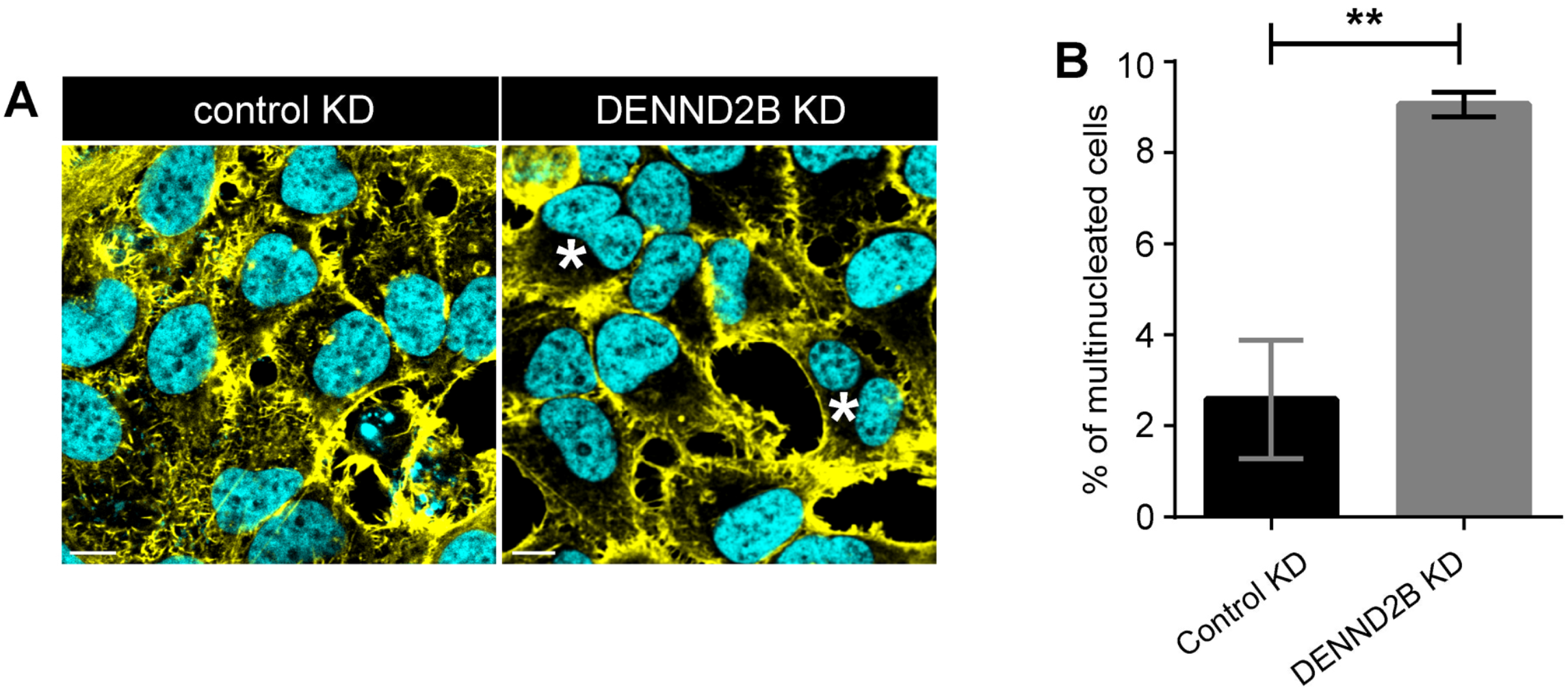
Loss of DENND2B increases number of binucleate cells. **(A)** HeLa cells were transduced with control or DENND2B shRNA lentivirus. Cells were fixed and stained with DAPI and Phalloidin. Scale bars = 10 μm **(B)** Quantification of experiment in (A); mean ± SEM; unpaired t-test (**, P ≤ 0.01; >60 cells per condition).

**Supplementary figure 2.**
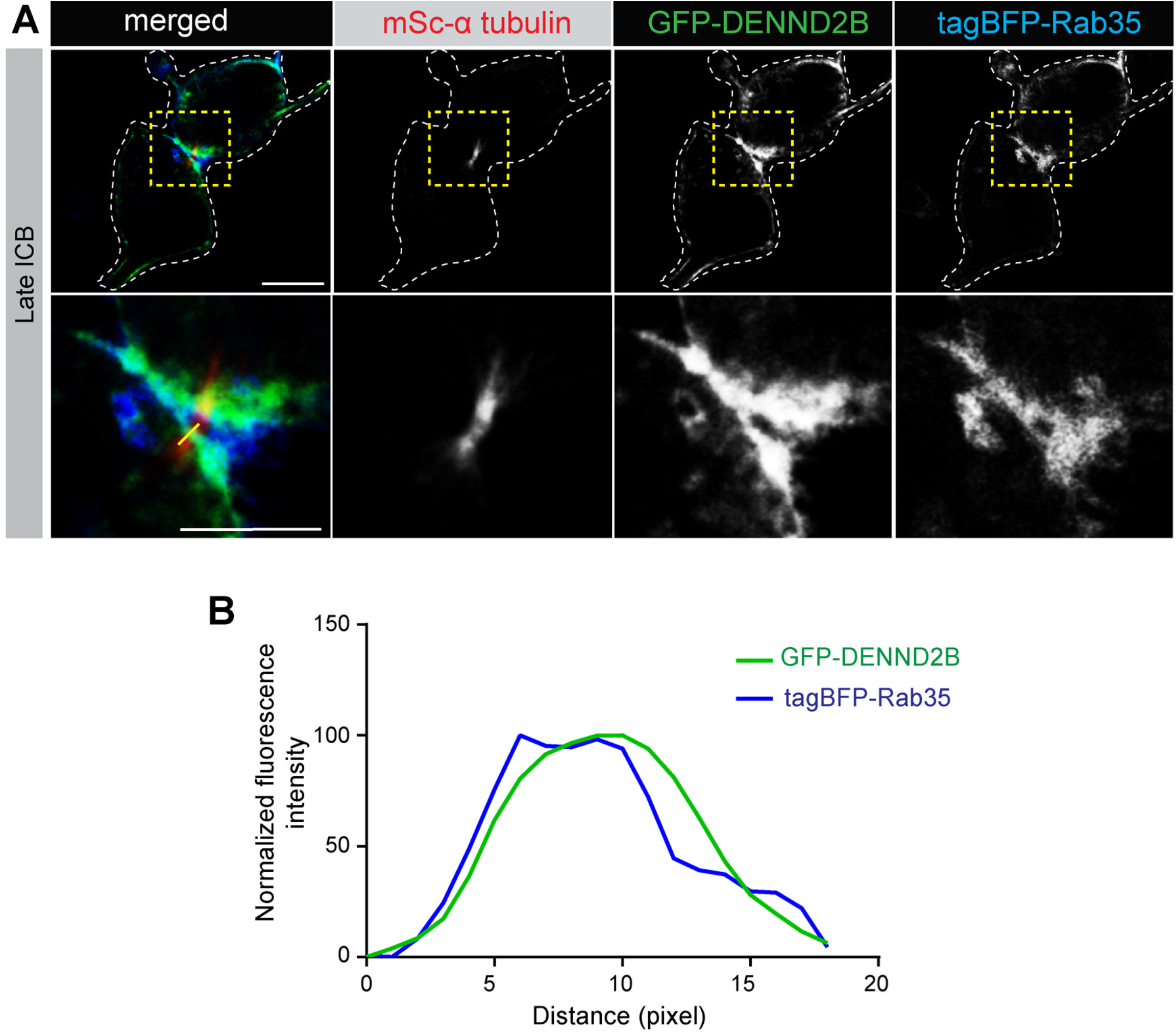
DENND2B and Rab35 colocalize at the cytokinetic bridge. **(A)** HeLa cells were transduced with lentiviruses mediating expression of GFP-DENDN2B, mSc-alpha tubulin and tagBFP-Rab35. Cells were fixed and stained with LAP2-β. Scale bars =13.2 μm for the low-magnification images and 6.25 μm for the higher magnification insets (represented with blue box). Dotted white line marks the periphery of dividing cell. **(B)** Normalized fluorescence intensity profiles along the yellow line from the inset image in (A) across the cytokinetic bridge.

**Supplementary figure 3.**
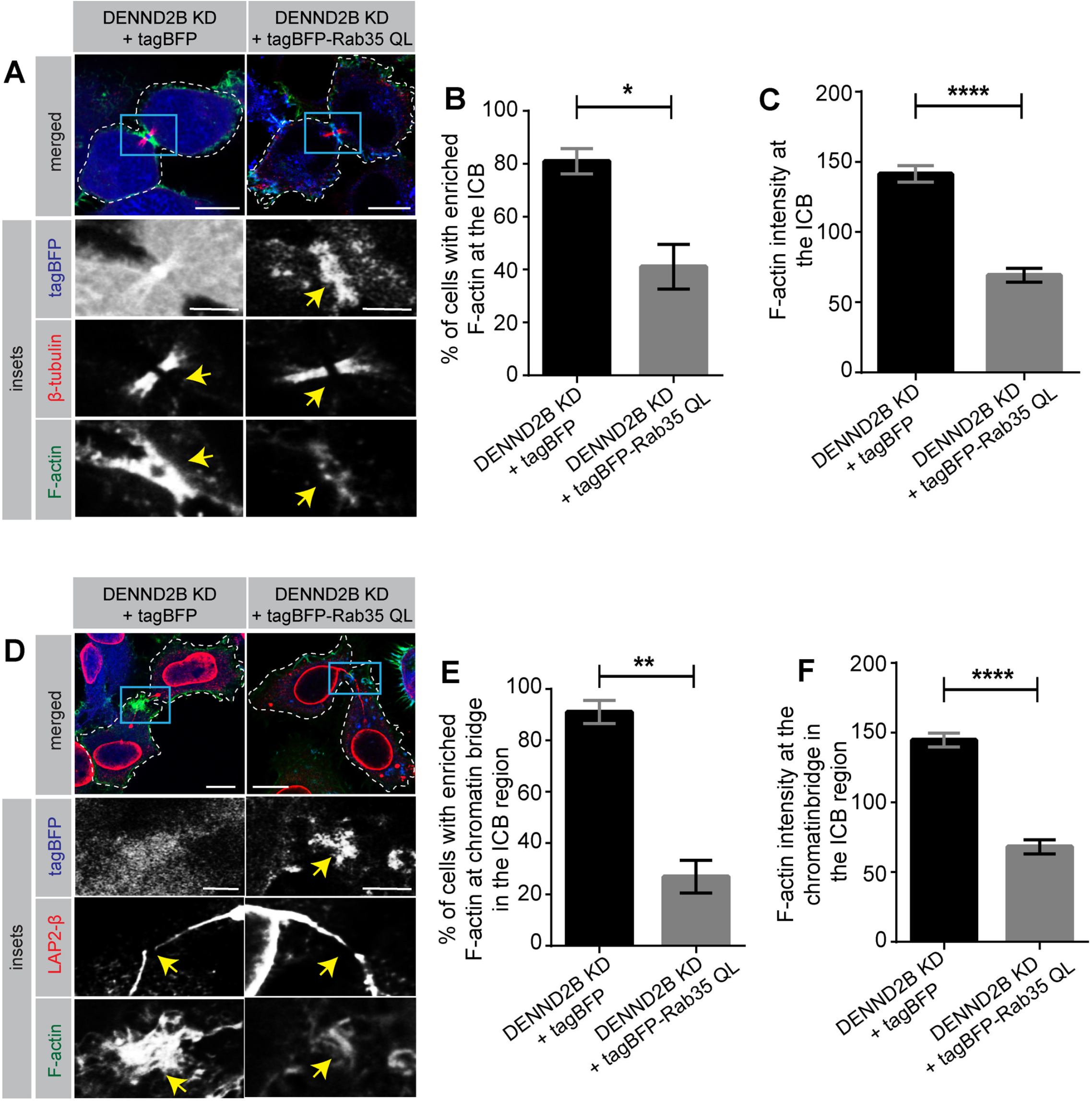
Expression of Rab35 active mutant rescues F-actin levels at the bridge with or without chromatin bridge. **(A)** HeLa cells transduced with DENND2B shRNA lentivirus were further transduced with lentivirus mediating expression of tagBFP and tagBFP-Rab35 QL. Cells were fixed and stained with β-tubulin and Phalloidin. Scale bars = 10 μm for the low-magnification images and 4.25 μm for the higher magnification insets (represented with blue box). Dotted white line marks the periphery of dividing cell. Yellow arrow represents the cytokinetic bridge. **(B)** Quantification of experiment in (A); mean ± SEM; unpaired t-test (*, P ≤ 0.05; >25 cells per condition). **(C)** Quantification of experiment in (A); mean ± SEM; Mann-Whitney U test (****, P ≤ 0.0001; >25 cells per condition). **(D)** HeLa cells transduced with DENND2B shRNA lentivirus were further transduced with lentivirus mediating expression of tagBFP and tagBFP-Rab35 QL. Cells were fixed and stained with LAP2-β and Phalloidin. Scale bars = 12 μm for the low-magnification images and 4.25 μm for the higher magnification insets (represented with blue box). Dotted white line marks the periphery of dividing cell. Yellow arrow represents the chromatin bridge. **(E)** Quantification of experiment in (C); mean ± SEM; unpaired t-test (**, P ≤ 0.01; >25 cells per condition). **(F)** Quantification of experiment in (C); mean ± SEM; Mann-Whitney U test (****, P ≤ 0.0001; >25 cells per condition).

